# Combined expression of FOXO and Tribbles proteins predicts survival of glioma patients

**DOI:** 10.1101/2025.01.20.633940

**Authors:** Bruno F. Santos, Ana-Teresa Maia, Inês Grenho, André Besouro-Duarte, Juan M Sepúlveda, Bibiana I. Ferreira, Wolfgang Link

**Author notes:** Correspondence to: Bibiana I. Ferreira., Wolfgang Link.

## Abstract

**Background:** Gliomas, particularly high-grade variants such as glioblastoma, remain therapeutically challenging with poor survival outcomes. Current biomarkers inadequately stratify patients for therapy selection, monitoring, and early detection of progression. The FOXO transcription factors and Tribbles pseudokinases, key regulators of the PI3K/AKT pathway, have been implicated in cancer progression but their prognostic value in gliomas is unclear. The aim of this study was to determine whether transcriptional profiles of FOXO and Tribbles family members predict survival in glioma patients better than established biomarkers.

**Methods:** Using RNA-seq data from The Cancer Genome Atlas (TCGA, **n** = 705) and Chinese Glioma Genome Atlas (CGGA, **n** = 1018), along with microarray datasets (REMBRANDT, n = 552; Gravendeel, n = 268), we analyzed mRNA levels of **FOXO1/3/4/6** and **TRIB1/2/3**. Survival analysis was performed via Kaplan-Meier curves, log-rank tests, and concordance indices. Gene expression differences across tumor grades and molecular subgroups were assessed using Student’s t-test.

**Results:** High *FOXO1* and low **FOXO3/4** mRNA levels, combined with elevated **TRIB1/2/3** expression, formed a signature significantly associated with worse survival across all cohorts. This signature stratifies grade 4 gliomas, with a median survival difference of 4.5 months. Grade 4 tumors exhibited elevated *FOXO1* and **TRIB1/2/3** but reduced **FOXO3/4** levels compared to lower-grade gliomas.

**Conclusions:** The FOXO/Tribbles transcriptional signature robustly predicts survival in glioma patients. These findings highlight its potential as a biomarker for patient stratification and a therapeutic target for high-grade gliomas.

**Key Points:**

1. FOXO/Tribbles mRNA signature predicts glioma survival status. Our analysis identifies a distinct transcriptional signature – characterized by high FOXO1 and TRIB1/2/3 levels along with low FOXO3 and FOXO4 – that consistently predicts poor prognosis across glioma cohorts. This FOXO/Tribbles signature demonstrates great prognostic power, particularly in grade 4 gliomas, providing an improved tool for risk stratification and clinical decision-making.
2. High FOXO1 and Tribbles levels mark aggressive glioblastoma tumors. We show that elevated expression of FOXO1 and Tribbles pseudokinases is enriched in glioblastoma (GB) and grade 4 astrocytomas. These tumors, known for their aggressiveness and therapy resistance, are consistently associated with this unfavorable mRNA profile. This suggests a functional role for FOXO1 and Tribbles in glioma progression and highlights them as promising biomarkers and therapeutic targets.
3. Multi-cohort validation (TCGA, CGGA) confirms prognostic robustness. The predictive FOXO/Tribbles expression pattern was validated across four large and independent datasets (TCGA, CGGA, REMBRANDT, and Gravendeel), demonstrating consistency despite differences in platform and population. This multi-cohort validation reinforces the robustness and generalizability of this transcriptional signature for predicting survival outcomes in glioma patients.

**Importance of the Study:** Glioblastoma and high-grade gliomas have dismal prognoses, yet current biomarkers fail to adequately guide therapy. This study identifies a novel transcriptional signature – combining FOXO transcription factors and Tribbles pseudokinases – that robustly predicts survival in glioma patients stratifying aggressive tumors. Using multi-cohort validation (TCGA, CGGA, REMBRANDT), we demonstrate that elevated *FOXO1* and Tribbles (**TRIB1/2/3**) levels, coupled with reduced **FOXO3/4**, correlate with poor outcomes in grade 4 gliomas. This signature is enriched in glioblastomas, highlighting its link to therapy resistance. Unlike prior studies focusing on individual proteins, our integrated analysis reveals their collective prognostic power, offering a pathway-specific biomarker for patient stratification. These findings provide a translational framework for targeting FOXO/Tribbles in precision oncology, addressing an urgent need for improved therapeutic strategies in neuro-oncology.

## Introduction

Glioblastoma (GB) is the most common and aggressive primary brain tumor in adults, accounting for almost half of the primary malignant brain tumors.^1^. In 2021, the World Health Organization (WHO) updated its classification of the central nervous system (CNS) tumors (WHO CNS 5^th^ edition classification, WHO CNS5) to incorporate molecular data, simplifying the system for improved diagnostic accuracy and prognostic assessment.^2^ GB treatment includes surgical resection followed by radiation therapy and Temozolomide (TMZ) chemotherapy. Nonetheless, GB remains an incurable disease with a median survival of only 15 months due to therapy resistance and recurrence.^3^ Therefore, there is an urgent need for new treatments that can overcome drug resistance, along with biomarkers to stratify risk or guide therapy.

Several studies revealed a prominent role of the PI3K/AKT signaling pathway in the formation and progression of gliomas, and it is possible to find changes in this pathway in approximately 90% of GB.^2,4^ We and others have characterized FOXO transcription factors as major transcriptional effectors of the PI3K/AKT pathway^5,6^ and identified the Tribbles homologue 2 (TRIB2) as a member of the Tribbles family of pseudokinases as a potent repressor of FOXO proteins^7^. Here we explore the role of FOXO and Tribbles proteins in driving aggressiveness in gliomas. FOXO proteins, a family of transcription factors including FOXO1, FOXO3, FOXO4 and FOXO6^8^, play a key role in regulating the expression of genes involved in cell proliferation, metabolism, apoptosis, autophagy, and stress resistance^9,10^ and are often inactivated in human cancers^11^. Additionally, variants of FOXO3 have been associated with extreme human longevity.^12^ A breakthrough in the functional characterization of FOXO transcription factors was achieved by the generation of triple knockout mice lacking FOXO1, 3 and 4, that established FOXO as *bona fide* tumor suppressors.^13^ Conversely, an oncogenic role of FOXOs in tumor progression, metastasis, and therapy resistance has emerged.^14^ These proteins are regulated by reversible post-translational modifications that convert external stimuli into specific transcriptional programs.^15^ Under stress conditions or in the absence of growth factors, FOXOs are active in the nucleus. Yet, in the presence of growth factors, or in cancer cells where the PI3K/AKT pathway is constitutively active, FOXOs are exported to the cytoplasm where they remain inactive.^10,16^ Thus, the pharmacological modulation of FOXO is considered an attractive therapeutic approach for treating cancer and age-related diseases.^17^ The extensive diversity of roles of these proteins suggests that the benefit of FOXO activation depends on the isoform and cellular context. While the different FOXO isoforms bind to the same consensus DNA sites in target gene promoters, it is increasingly evident that they also exert non-redundant functions in specific tissues.^18^ The Tribbles protein family consists of three isoforms, TRIB1, TRIB2 and TRIB3. Instead of phosphorylating target proteins, the members of this family function as protein scaffolds that integrate and modulate signals of important signaling pathways involved in proliferation, survival, and differentiation.^19^ Our group established TRIB2 as an oncogene in melanoma and found that high levels of this protein confer resistance to several drugs used for cancer treatment.^7,20,21^ While TRIB2 has been implicated in cancer progression and therapy resistance across numerous cancer types^22^, and TRIB1 is also considered as an oncogene in various malignancies, the role of TRIB3 is more complex and context-dependent. Studies indicate that TRIB3’s contribution to cancer progression varies, potentially acting as either a tumor suppressor or an oncogene depending on the specific cellular environment.^23–27^ However, the clinical relevance of these observations has yet to be fully established.

Here, we reveal a relationship between survival and mRNA levels of FOXO and Tribbles family members in patients with gliomas. We show that specific FOXO/Tribbles signatures are highly predictive for clinical outcome, with high FOXO1 and low FOXO3/4 plus high TRIB1/2/3 levels associated with a worse prognosis. Furthermore, this signature could be used to select patients with glioma who would benefit from new potential therapeutic targets.

## Materials and Methods

### Datasets

#### RNA-seq Datasets

RNA-seq data was obtained from two independent databases, The Cancer Genome Atlas (TCGA, https://portal.gdc.cancer.gov/) and The Chinese Glioma Genome Atlas (CGGA, http://www.cgga.org.cn/).

Two independent datasets (TCGA-GBM and TCGA-LGG), comprising a total of 705 samples, were directly downloaded from the TCGA database into RStudio (version 2023.3.0.386) in September 2024 using the TCGAbiolinks R package.^28^ TCGA data was reclassified according to the WHO CNS5 standards.^29^ The reclassified dataset comprises 113 grade 2 astrocytomas, 98 grade 3 astrocytomas, 24 grade 4 astrocytomas, 81 grade 2 oligodendrogliomas, 70 grade 3 oligodendrogliomas, 221 GB and 98 unclassified samples.

Data from the CGGA database (1018 samples) was obtained from two independent datasets (mRNAseq_693 and mRNAseq_325), downloaded from the CGGA portal in September 2024 and uploaded to RStudio. These datasets comprise 175 grade 2 astrocytomas, 214 grade 3 astrocytomas, 112 grade 2 oligodendrogliomas, 9 grade 3 oligodendrogliomas, 94 grade 2 mixed type, 21 grade 3 mixed type, 388 GB and 5 unclassified samples.

For both TCGA and CGGA datasets, read counts (STAR-counts) were converted to transcript per million (TPM) to reduce technical variability between datasets. Suppl. Table 1 provides an overview of the demographic characteristics and disease features for both RNA-seq datasets. Expression levels of FOXOs and Tribbles genes were analyzed by plotting Log2 transformed TPM values using the ggplot2 R package.^30^ Boxplots were compared using the Student’s t-test.

#### Microarray datasets

REMBRANDT (GSE68848) and the Gravendeel (GSE16011) microarray cohorts (Affymetrix HG-U1333Plus2) were available from Gene Expression Omnibus (GEO, https://www.ncbi.nlm.nih.gov/geo/query/acc.cgi). Both databases were downloaded in September 2024 and uploaded into RStudio using GEOquery^31^ and Biobase^32^ R packages. In cases where multiple probes were used for one gene, their average expression levels were used. Matching clinical data for REMBRANDT and Gravendeel datasets was obtained through GEO or GlioVis (https://gliovis.bioinfo.cnio.es/), respectively. Rembrandt dataset comprises 61 grade 2 astrocytomas, 53 grade 3 astrocytomas, 27 grade 2 oligodendrogliomas, 22 grade 3 oligodendrogliomas, 4 grade 2 mixed type, 3 grade 3 mixed type, 190 GB and 192 unclassified samples. The Gravendeel dataset comprises 13 grade 2 astrocytomas, 16 grade 3 astrocytomas, 8 grade 2 oligodendrogliomas, 44 grade 3 oligodendrogliomas, 3 grade 2 mixed type, 25 grade 3 mixed type and 159 GB. Suppl. Table 1 provides an overview of the demographic characteristics and disease features for both microarray datasets.

### Survival Analysis

Survival analysis was performed with Kaplan-Meier, using the survival^33^ and survminer^34^ R packages. The primary endpoint was death from any cause, with surviving subjects censored at the date of their last known contact. Kaplan–Meier survival curves were compared using the log-rank test. To quantify the correlation between risk predictions and event times, maximizing the ability to better discriminate patient survival, their Concordance Index was determined using the concordance function from survival R package.^33^

### Cox Proportional Hazards Analysis

Multivariate Cox proportional hazards regression was performed using the survival^33^ R package to assess the independent prognostic value of the combined transcriptional profiles of FOXO and Tribbles on the overall survival. Hazard ratios (HRs) and 95% confidence intervals (CIs) were estimated by fitting the Cox model while adjusting for age (stratified into young, middle-aged, and elderly^35^), gender, IDH mutation status, 1p/19q codeletion, MGMT promoter methylation, and WHO classification.

### Ethics Statement

This study involved only the analysis of publicly available, de-identified human gene expression and clinical data from The Cancer Genome Atlas (TCGA), the Chinese Glioma Genome Atlas (CGGA), the REMBRANDT dataset (GSE68848), and the Gravendeel dataset (GSE16011). As such, no institutional review board approval or informed consent was required. All data used are compliant with relevant ethical guidelines and regulations.

### Code availability

The filtered data and code for the survival analysis of gliomas according to FOXOs and Tribbles expression levels can be publicly accessed at https://github.com/maialab/enduring.

### FOXO1 and TRIB3 depletion

Gene depletion was achieved through siRNA-mediated silencing for FOXO1 and CRISPR/Cas9-mediated knockout for TRIB3. U87 glioblastoma cells were transfected with siRNA targeting FOXO1 (25 nM; Dharmacon™, L-003006-00-0005) or a non-targeting scramble control (25 nM; Dharmacon™, LD-001810-01-20) using Dharmacon™ DharmaFECT™ siRNA Transfection Reagent according to the manufacturer’s protocol. Cells were harvested 72 hours post-transfection for protein extraction and immunoblot analysis to confirm FOXO1 knockdown. Parallel cultures were used for functional assays, including cell viability measurements. For TRIB3 depletion, U118 glioblastoma cells were edited using the CRISPR/Cas9 system. Two guide RNAs targeting exon one of TRIB3 (sequence: 5’-GCCCACTTCGAGCTCGTTTC-3’; 5’-GTTGCACGATCTGGAGCAGT-3’) were cloned into a Cas9 expression vector construct (Plasmid *#62988*, Addgene) and delivered into cells via Lipofectamine 2000 transfection. Following antibiotic selection (1μg/ul Puromycin), single-cell clones were isolated by limiting dilution and expanded. Successful TRIB3 knockout was verified by immunoblotting, with wild-type U118 cells maintained in parallel as controls.

### Functional assays

Cell viability was assessed using the MTT assay. Briefly, U118 wild-type and TRIB3 knockout cells, as well as U87 cells transfected with either FOXO1-targeting siRNA or non-targeting scramble control, were seeded in 96-well plates at a density of 5×10^3^ cells per well in complete growth medium. After 72 hours, cells were incubated with MTT solution (0,5 mg/mL; Alfa Aesar, L11939) for 3 hours at 37°C. Formazan crystals were dissolved by adding 100 µL of DMSO, and absorbance was measured at 570 nm using a microplate reader. Cell viability was calculated as a fold change relative to wild-type cells (for U118) or scramble-transfected cells (for U87). Experiments were performed in triplicate and repeated at least three independent times.

For protein validation, cells were lysed in RIPA buffer (Tris pH 7.5 (20mM) (Fisher Scientific, USA), sodium chloride (NaCl) (150mM) (Merck, Germany), 1% Triton X-100 (Amresco, USA), NaF (50mM) (VWR, USA), ethylenediaminetetraacetic acid (EDTA) (1mM) (Sigma Aldrich, USA), aminopolycarboxylic acid (EGTA) (1mM) (AppliChem, Germany), pyrophosphate (Santa Cruz, USA) (2,5mM), β-glycerophosphate (b-g-p) (2mM) (Santa Cruz, USA), OVO4 (1mM) (Sigma Aldrich, USA), calyculin A (100nM) (Santa Cruz, USA) and 1x protease inhibitors cocktail (Thermo Fisher, USA) and protein concentrations were determined by BCA assay. Equal amounts of protein were separated by SDS-PAGE, transferred to PVDF membranes, and blocked with 5% non-fat milk in TBS-T. Membranes were incubated with primary antibodies against TRIB3 (AB75846, Abcam) or FOXO1 (#2880, CST), followed by HRP-conjugated secondary antibodies. Bands were visualized using chemiluminescence, and GAPDH (SC-25778, Santa Cruz) or Tubulin (T9026, Sigma) served as loading controls.

## Results

### High FOXO1, low FOXO3/FOXO4, and elevated TRIB1/2/3 mRNA levels form a molecular signature predictive of clinical outcomes

To investigate if mRNA levels of FOXO family members were associated with the prognosis of glioma patients, we analyzed the mRNA levels of a total of 705 samples (534 Low-Grade Gliomas (LGG) and 171 GB) from the TCGA database, using RStudio. Quartiles were established as the cut-off to define low expression levels under the first quartile and high levels above the third quartile. Kaplan-Meier curves show that the patient survival rate is dependent on FOXO mRNA levels. High levels of FOXO1 (p<0.0001, Fig. 1A) and low levels of FOXO3 (p<0.0001, Fig. 1B), FOXO4 (p<0.0001, Fig. 1C) or FOXO6 (p=0.027, Fig. 1D) are significantly associated with a worse prognosis. Interestingly, FOXO6 mRNA levels have a considerably lower capacity to separate patients according to survival than the rest of FOXOs (Fig. 1D). Given that FOXO expression is tissue-specific, and their functions can overlap, with one isoform potentially compensating for another, we assessed whether the combined expression of various FOXO isoforms was associated with patient prognosis. We observed that patients with poorer survival rates share the characteristic of having high mRNA levels of FOXO1, along with low levels of FOXO3 and FOXO4, irrespective of their FOXO6 levels (Suppl. Fig. 1A). We validated our findings using an independent dataset of glioma RNA-seq data (CGGA), comprising 1018 samples, and found that high expression of FOXO1 (p<0.0001, Suppl. Fig. 1B) or low expression of FOXO4 (p<0.0001, Suppl. Fig. 1D) was significantly associated with poorer patient survival. In contrast, transcript levels of FOXO3 (p=0.25, Suppl. Fig. 1C) and FOXO6 (p=0.45, Suppl. Fig. 1E) did not show a significant correlation with patient survival.

**Fig. 1.**
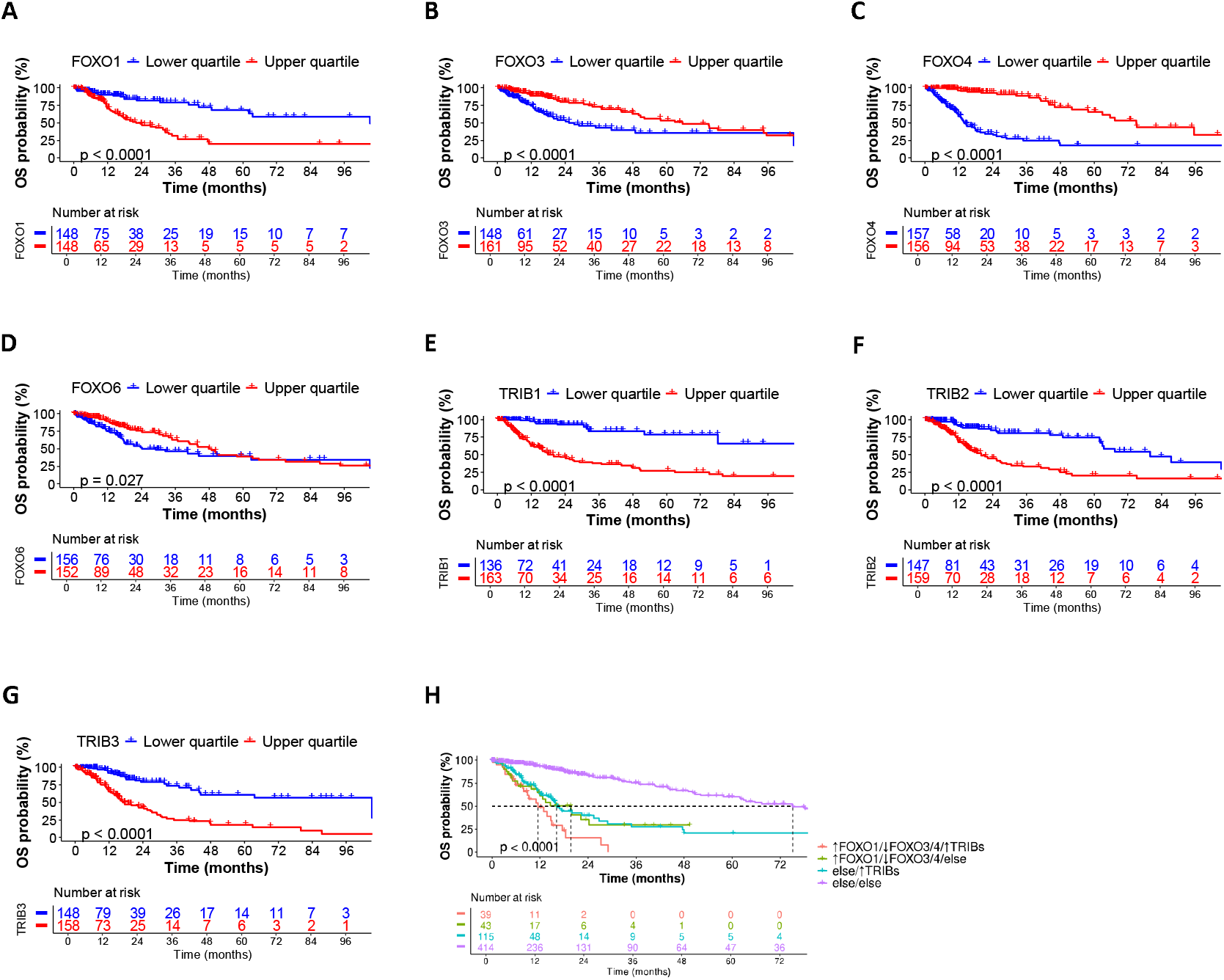
High FOXO1, low FOXO3/FOXO4, and elevated TRIB1/2/3 mRNA levels form a molecular signature predictive of clinical outcomes. **A-G:** Kaplan-Meier plots of FOXO and Tribbles for TCGA glioma samples. The lower quartile, in blue, and the upper quartile, in red, were established as the cut-off for low and high expression levels, respectively. **A:** FOXO1; **B:** FOXO3; **C:** FOXO4; **D:** FOXO6; **E:** TRIB1; **F:** TRIB2; **G:** TRIB3. **H:** Kaplan-Meier plot of a combination of high levels of FOXO1 and low levels of FOXO3/4 plus high levels of Tribbles for TCGA glioma samples. The median value was set as the cut-off. “Else” means every other possible combination of expressions than the stated in comparison. High FOXO1 and low FOXO3/4 mRNA levels combined with high levels of TRIB1/2/3 (↑FOXO1/↓FOXO3/4/↑TRIBs) are represented in pink. All p-values refer to log-rank test.

Next, we sought to investigate if Tribbles transcript levels would also be associated with different survival rates. We found that high mRNA levels of TRIB1 (p<0.0001, Fig. 1E), TRIB2 (p<0.0001, Fig. 1F) or TRIB3 (p<0.0001, Fig. 1G) were significantly associated with worse patient prognosis. Moreover, patients who expressed combined high levels of all three Tribbles members were associated with the worst survival rates (Suppl. Fig. 1F). Importantly, elevated mRNA expression of TRIB1 (p<0.0001, Suppl. Fig. 1G), TRIB2 (p<0.0001, Suppl. Fig. 1H), or TRIB3 (p<0.0001, Suppl. Fig. 1I) in CGGA patients was associated with significantly worse survival, aligning with our results from the TCGA dataset.

FOXO transcription factors are downstream effectors of the PI3K/AKT pathway whose activity is modulated by TRIB2 and/or TRIB3 levels in many cancers.^19^ Therefore, we sought to investigate if combined expression of these proteins would have a major impact on prognosis in gliomas. Our analysis revealed that patients who simultaneously expressed high levels of FOXO1 and low levels of FOXO3/4 plus high levels of TRIB1/2/3 represent the worst prognosis group of gliomas patients, for both TCGA (p<0.0001, Fig. 1H) and CGGA (p<0.0001, Suppl. Fig. 1J) datasets. These findings suggest a potential functional interplay between FOXO isoforms and Tribbles family proteins in promoting glioma progression and highlight this specific molecular signature as a powerful prognostic indicator for poor clinical outcomes in glioma patients.

### Independent glioma datasets validate the mRNA signature and its robustness

To further validate our findings, we analyzed two independent microarray datasets: REMBRANDT (GSE68848) and Gravendeel (GSE16011). In the REMBRANDT dataset, which includes 552 glioma samples, the expression patterns of FOXO family members showed to be consistent with the predictive signature described above. Specifically, elevated FOXO1 expression levels (p=0.032, Fig. 2A) and reduced levels of FOXO3 (p<0.0001, Fig. 2B), FOXO4 (p=0.00065, Fig. 2C), and FOXO6 (p=0.00042, Fig. 2D) were significantly associated with poorer patient prognosis. Regarding the Tribbles family members, high expression levels of TRIB2 (p=0.0091, Fig. 2F) and TRIB3 (p<0.0001, Fig. 2G) correlated with worse survival rates. However, no significant association was found between TRIB1 transcript levels and patient survival (p=<0.25, Fig. 2E). The combined expression of these proteins with the worst impact on prognosis revealed to be in the patients who simultaneously expressed low levels of FOXO1/3/4/6 plus high levels of TRIB1/2/3 (↓FOXOs/↑TRIBs) (p<0.0001, Fig. 2H).

**Fig. 2.**
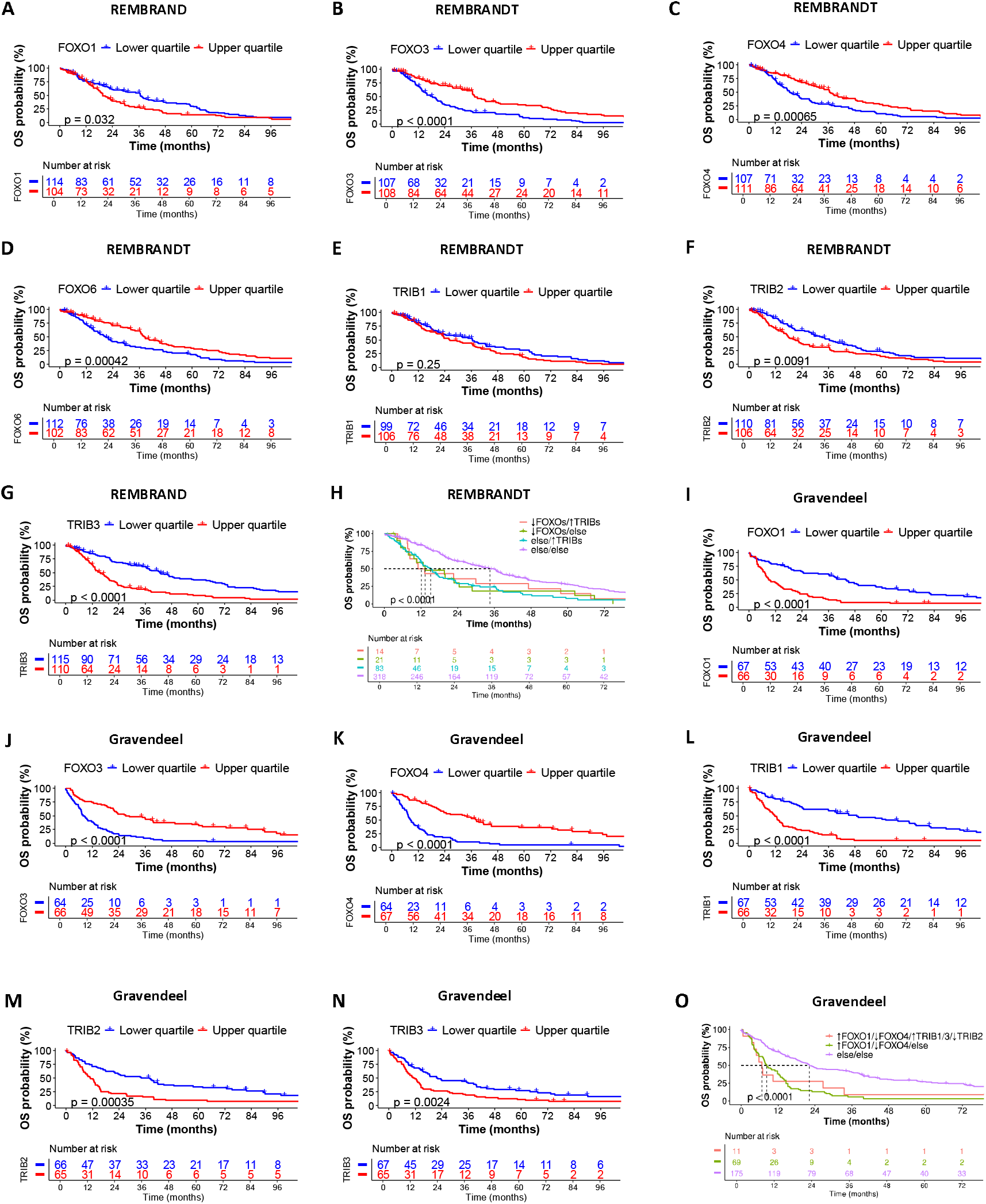
REMBRANDT and Gravendeel microarray datasets validate the mRNA signature and its robustness. **A-G:** Kaplan-Meier plots of FOXO and Tribbles for REMBRANDT glioma samples. The lower quartile, in blue, and the upper quartile, in red, were established as the cut-off for low and high expression levels, respectively. **A:** FOXO1; **B:** FOXO3; **C:** FOXO4; **D:** FOXO6; **E:** TRIB1; **F:** TRIB2; **G:** TRIB3. All p-values refer to log-rank test. **H:** Kaplan-Meier plot of a combination of low levels of FOXOs plus high levels of Tribbles for REMBRANDT glioma samples. The median value was set as the cut-off. “Else” means every other possible combination of expressions than the stated in comparison. Low FOXO1/3/4/6 mRNA levels combined with high levels of TRIB1/2/3 (↓FOXOs/↑TRIBs) are represented in pink. All p-values refer to log-rank test. **I-N:** Kaplan-Meier plots of FOXO and Tribbles for Gravendeel glioma samples. The lower quartile, in blue, and the upper quartile, in red, were established as the cut-off for low and high expression levels, respectively. **I:** FOXO1; **J:** FOXO3; **K:** FOXO4; **L:** TRIB1; **M:** TRIB2; **N:** TRIB3. All p-values refer to log-rank test. **O:** Kaplan-Meier plot of a combination of high levels of FOXO1 and low levels of FOXO4 plus high levels of TRIB1/3 and low levels of TRIB2 for Gravendeel glioma samples. The median value was set as the cut-off. “Else” means every other possible combination of expressions than the stated in comparison High FOXO1 and low FOXO4 mRNA levels combined with high levels of TRIB1/3 and low of TRIB2 (↑FOXO1/↓FOXO4/↑TRIB1/3/↓TRIB2) are represented in pink. All p-values refer to log-rank test.

In the Gravendeel microarray dataset, which includes 268 glioma samples, the association between expression levels and survival was consistent with our findings from the TCGA dataset. Specifically, low levels of FOXO3 (p<0.0001, Fig. 2J) and FOXO4 (p<0.0001, Fig. 2K), along with high levels of FOXO1 (p<0.0001, Fig. 2I), TRIB1 (p<0.0001, Fig. 2L), TRIB2 (p=0.00035, Fig. 2M) and TRIB3 (p=0.0024, Fig. 2N) are significantly correlated with poorer prognosis. Notably, this dataset did not include expression levels for FOXO6. For these samples, high levels of FOXO1 and low levels of FOXO4 plus high levels of TRIB1/3 and low levels of TRIB2 (↑FOXO1/↓FOXO4/↑TRIB1/3/↓TRIB2) represent the worst prognosis group of glioma patients (p<0.0001, Fig. 2O).

Taken together, these findings show that, despite some variability across different populations, patients with low FOXO3, FOXO4 and FOXO6, and with high FOXO1, TRIB1, TRIB2 and TRIB3 mRNA levels have significantly worse survival rates. Discrepancies among FOXO3, FOXO6 and TRIB1 suggest that further studies are needed to fully establish their relationship with glioma patient survival. Importantly, the data suggests that combined transcriptional profiles of FOXO and Tribbles family members could be used to predict the clinical outcome of gliomas patients.

### High grade tumors express ↑ FOXO1/↓FOXO3/4/↑TRIB1/2/3 mRNA levels

Next, we sought to determine whether certain glioma tumor types were overrepresented among those exhibiting a high FOXO1 and high Tribbles expression levels accompanied by low FOXO3/4 expression levels. We found that the ↑FOXO1/↓FOXO3/4/↑TRIBs expression pattern was exclusively observed in patients diagnosed with glioblastoma (GB), or grade 4 astrocytoma (Fig. 3A and 3B). In fact, the majority of GB cases exhibited either the ↑FOXO1/↓FOXO3/4 or ↑TRIBs profile, or both. This finding suggests that this particular mRNA expression signature is strongly associated with higher-grade gliomas known to be associated with poorer prognosis and diminished overall survival (Suppl. Fig. 2A). Consistently, we hypothesized that the mRNA levels of FOXO1, FOXO3, FOXO4, TRIB1, TRIB2, and TRIB3 varied according to tumor grade. To address this question, we evaluated the mRNA expression levels, from the TCGA database, of each individual FOXO or Tribbles gene between GB and other gliomas. We found that FOXO1 (p<0.0001, Fig. 3C), TRIB1 (p<0.0001, Fig. 3F), TRIB2 (p<0.0001, Fig. 3G) and TRIB3 (p<0.0001, Fig. 3H) are overexpressed in GBs compared to patients with other type of gliomas, whilst FOXO3 (p<0.0001, Fig. 3D) and FOXO4 (p<0.0001, Fig. 3E) mRNA levels are underexpressed.

**Fig. 3.**
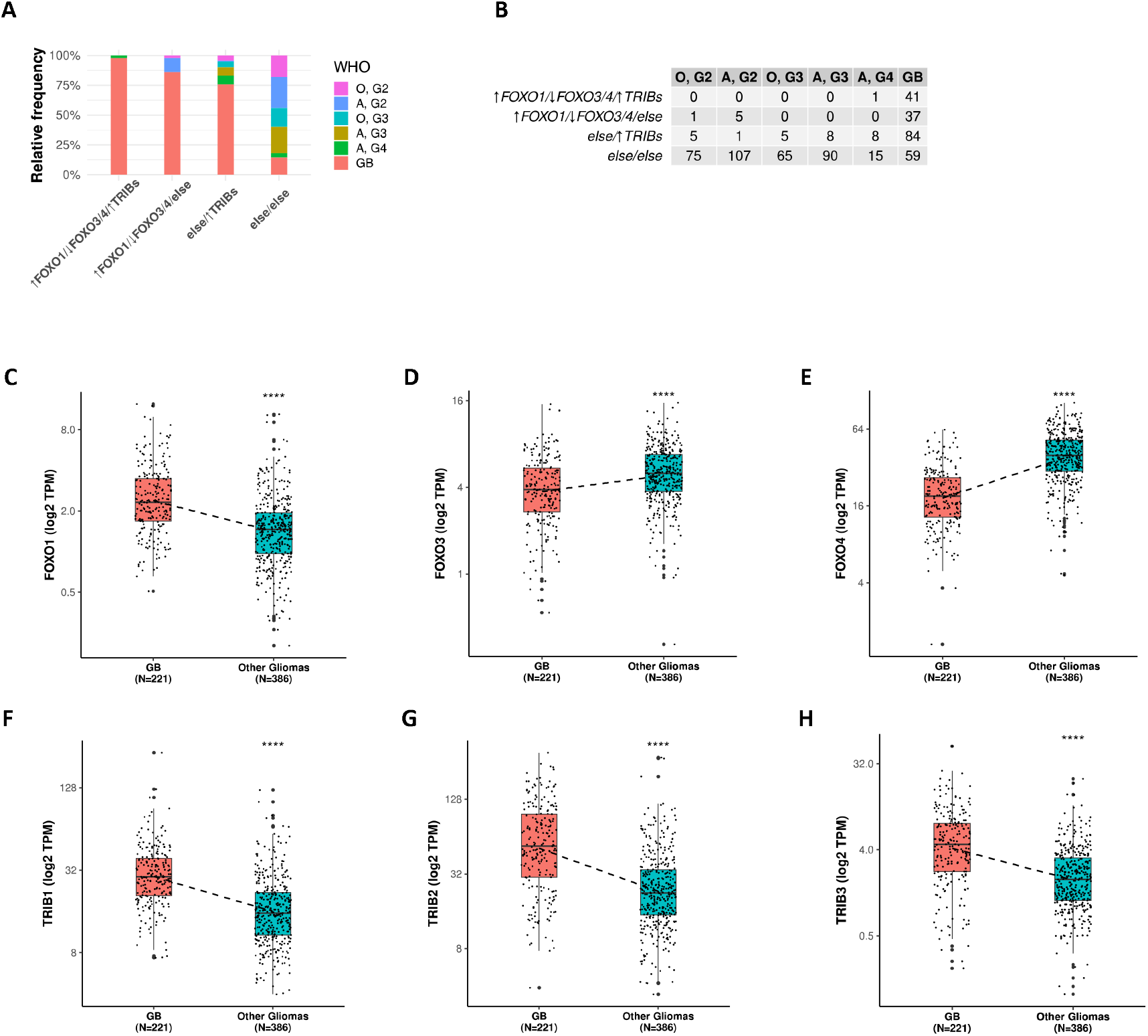
High grade tumors express ↑ FOXO1/↓FOXO3/4/↑TRIB1/2/3 mRNA levels. **A:** Percentage of the different types of tumors for each combination of high levels of FOXO1 and low of FOXO3/4 (↑FOXO1/↓FOXO3/4) and high levels of Tribbles (↑TRIBs) for TCGA glioma samples. “Else” means every other possible combination of expressions than the stated in comparison. **B:** Number of cases of the different types of tumors for each combination represented in A. **C-H:** Boxplots of FOXO and Tribbles mRNA levels among glioblastoma and other glioma types of TCGA samples. N represents the number of samples in each group. **C:** FOXO1; **D:** FOXO3; **E:** FOXO4; **F:** TRIB1; **G:** TRIB2; **H:** TRIB3. All p-values refers to Student’s t-test: **** p<0.0001

To evaluate the independent prognostic impact of the combined transcriptional profiles of FOXO and Tribbles, we performed a multivariate Cox proportional hazards model, adjusted for established clinical and molecular variables (Fig. 4 and Suppl. Table 2). The model included age group, gender, IDH mutation status, 1p/19q codeletion, *MGMT* promoter methylation, WHO classification, and FOXO/Tribbles expression profiles. Age was a strong predictor of overall survival, with elderly patients (>64 years) showing a significantly higher risk of death compared with the youngest group (<48 years) (HR = 5.30, 95%, p = 3.5 × 10^−8^), while middle-aged patients (48–64 years) had a moderate increase (HR = 1.69, p = 0.049). Among molecular markers, wild-type IDH (HR = 3.27, p = 0.020) and non-codeleted 1p/19q status (HR = 14.06, p < 0.001) were associated with poorer survival, whereas MGMT methylation showed no significant effect (p = 0.42). Tumor classification was also prognostically relevant: Oligodendroglioma Grade 3 cases exhibited increased risk compared with Grade 2 (HR = 4.07, p = 0.019), while Astrocytoma Grades 2 and 3 were associated with a significantly lower hazard (HR = 0.19, p = 0.001; HR = 0.35, p = 0.029, respectively). Importantly, the analysis revealed no significant differences in survival between Astrocytoma Grades 4 and GB, showing that they behave similarly in prognostic terms. Although the FOXO/Tribbles molecular profiles did not reach statistical significance, this multivariate model confirms that the proposed signature retains independent prognostic value when adjusted for these established clinical and molecular factors, particularly the ↑FOXO1/↓FOXO3/4 or ↑TRIBs profile (HR = 1.35, p = 0.33). Absence of statistical significance may indicate that their prognostic effect may be influenced by overlapping clinical or molecular variables.

**Fig. 4.**
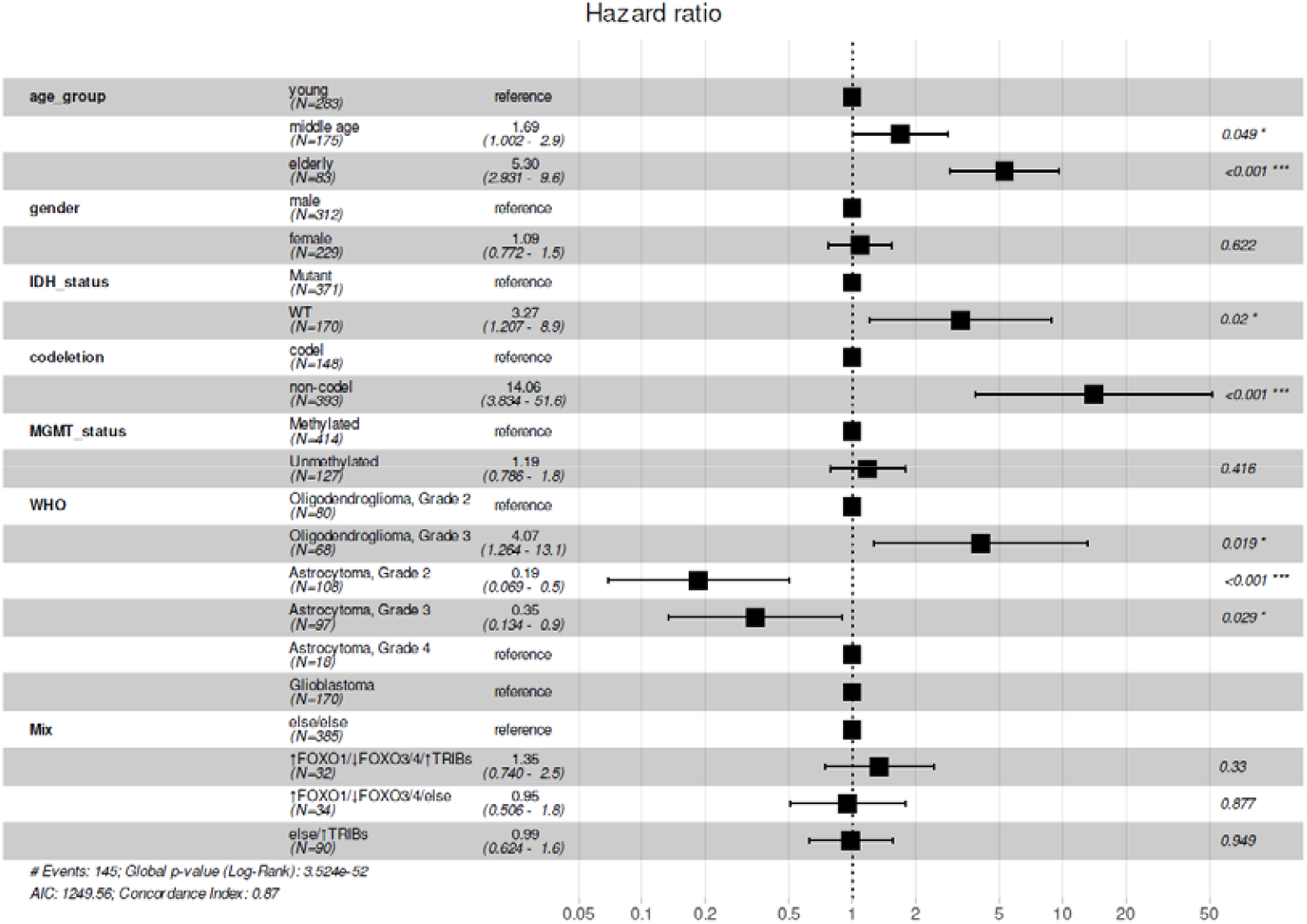
Multivariate Cox proportional hazards regression model assessing potential confounding factors for the combined transcriptional profiles of FOXO and Tribbles. The analysis included age group, gender, *IDH* mutation status, 1p/19q codeletion, *MGMT* promoter methylation, WHO classification, and FOXO/Tribbles expression profiles as covariates. Hazard ratios (HR) with corresponding 95% confidence intervals (CI) and p-values were estimated using a Cox proportional hazards model.

### FOXOs and Tribbles expression enable enhanced stratification of grade 4 glioma patients

The presence of the IDH mutation in gliomas is a favorable prognostic marker (p<0.0001,Fig. 5A) and targeted therapeutics have been approved to treat patients with grade 2 astrocytoma or oligodendroglioma with mutated IDH. However, patients without mutations in the IDH genes (i.e. patients with GB) are of greater concern as they exhibit worse survival rates and face a lack of effective treatment options. Therefore, it is paramount to find new biomarkers that further stratify these patients and ultimately develop new drugs capable of targeting specific drivers of these tumors. We hypothesized that a signature of high FOXO1, low FOXO3/4, and high TRIB1/2/3 could identify patients likely to benefit from novel therapeutics targeting these families, which are currently in preclinical development. The majority of patients with high FOXO1 and low FOXO3/4 signature, elevated TRIB1/2/3 levels, or both signatures combined, are IDH wild-type (Fig. 5B). This is unsurprising since these signatures are enriched in GB (Fig. 3A). These findings suggest patients that lack viable treatment options may benefit from evaluating FOXO and Tribbles protein levels as potential therapeutic markers

**Fig. 5.**
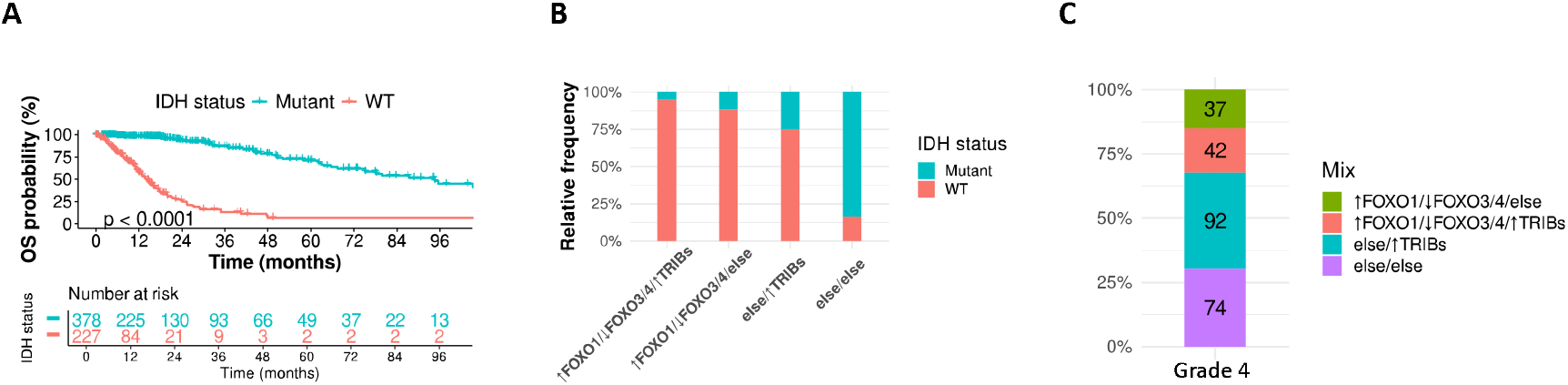
FOXOs and Tribbles expression enable enhanced stratification of grade 4 glioma patients. **A:** Kaplan-Meier plot of IDH mutation status for the TCGA glioma samples. Mutated in pink and wild type (wt) in blue. **B:** Percentage of patients with different IDH mutation status for each combination of high levels of FOXO1 and low of FOXO3/4 (↑FOXO1/↓FOXO3/4) and high levels of Tribbles (↑TRIBs) for TCGA glioma samples. “Else” means every other possible combination of expressions than the stated in comparison. **C:** Number of patients for each combination of high FOXO1 and low FOXO3/4 levels plus high levels of Tribbles for TCGA grade 4 glioma samples. **D:** Kaplan-Meier plot of MGMT promoter methylation status for the TCGA grade 4 glioma samples. Methylated in pink and unmethylated in blue. **E:** Kaplan-Meier plot of a combination of FOXOs and Tribbles levels in TCGA grade 4 glioma samples. Subgroups with high FOXO1 and low FOXO3/4 mRNA levels (↑FOXO1/↓FOXO3/4/else), high levels of TRIB1/2/3 (else/↑TRIBs), or both (↑FOXO1/↓FOXO3/4/↑TRIBs) represented together in pink. “Else” means every other possible combination of expressions than the stated in comparison. All p-values refer to log-rank test.

Accordingly, we aimed to analyze the robustness of the FOXO/Tribbles signature in stratifying GB patients, as well as grade 4 Astrocytomas. Given the aggressive nature of these high-grade tumors both patient groups typically receive similar treatment regimens, making this comparison particularly relevant.^40^ We first analyzed the distribution of TCGA patients with grade 4 tumors (astrocytomas or glioblastomas) across different combinations of FOXO and Tribbles signatures. We found that approximately 70% of grade 4 glioma patients express high levels of TRIB1/2/3 and/or decreased FOXO3/4 levels compared to 30% of patients who show no alteration on mRNA levels of both TRIBBLEs and FOXO (Fig. 5C). This means that the majority of grade 4 patients, approximately 55%, would potentially benefit from Tribbles inhibition while 32% could benefit from FOXO modulation. Of these patients eligible for target therapy, 25% would profit from both FOXO and Tribbles therapeutics.

We next compared the discriminatory power of the FOXO/Tribbles signature to that of established glioma biomarkers. A key limitation of the available datasets is the lack of detailed treatment information, particularly on the use and type of chemotherapy. This gap impedes our ability to distinguish whether the observed survival benefits are prognostic or predictive. For instance, while MGMT promoter methylation status is an established predictive biomarker in GB, associated with improved response to temozolomide, its independent prognostic value remains debated.^41,42^ Likewise, in astrocytomas *MGMT* promoter is frequently methylated, although its clinical relevance has not yet been as well established as in GB.^2^ In order to evaluate the potential clinical relevance of the FOXO/Tribbles signature, we compared its performance to *MGMT* promoter methylation using Kaplan–Meier analysis and calculated the Concordance Index, a metric of the discriminatory ability of a prognostic model (with higher values indicating better discrimination). For the MGMT status, TCGA patients with grade 4 were separated into methylated and unmethylated (p=0.018, Fig. 5D). For the FOXO/Tribbles signature, TCGA patients with grade 4 were separated into patients hypothetically eligible for FOXO and/or Tribbles therapeutics against else/else subgroup of patients (p=0.023, Fig. 5E, Suppl. Fig. 3).

Our results show that the FOXO/Tribbles signatures offered prognostic discrimination of grade 4 patients in terms of survival beyond epigenetic inhibition of *MGMT*. High FOXO1 and low FOXO3/4 signature, high levels of TRIB1/2/3, or both signatures combined, collectively show a median survival difference of 4.5 months compared to other signature combinations, with a Concordance Index of 0.55. This contrasts with the 2.7-month difference observed for MGMT status (methylated vs. unmethylated), with a Concordance Index of 0.46. These findings suggest that, independently of MGMT status, the FOXO/Tribbles signature has meaningful discriminatory power that may support its utility for patient stratification in the clinical setting.

Based on the contentious nature of FOXO1 and TRIB3 in cancer biology, coupled with our data demonstrating their clear oncogenic roles, we sought to experimentally validate these findings in GB cells. To this end, we depleted FOXO1 and TRIB3 from U87 and U118 GB cells using RNAi mediated silencing or CRISPR/Cas9-mediated knockout, respectively. Figure 6 shows that loss of either TRIB3 or FOXO1 reduced GB cell viability, suggesting that FOXO1 and TRIB3 expression support survival pathways in glioblastoma cells.

**Fig. 6.**
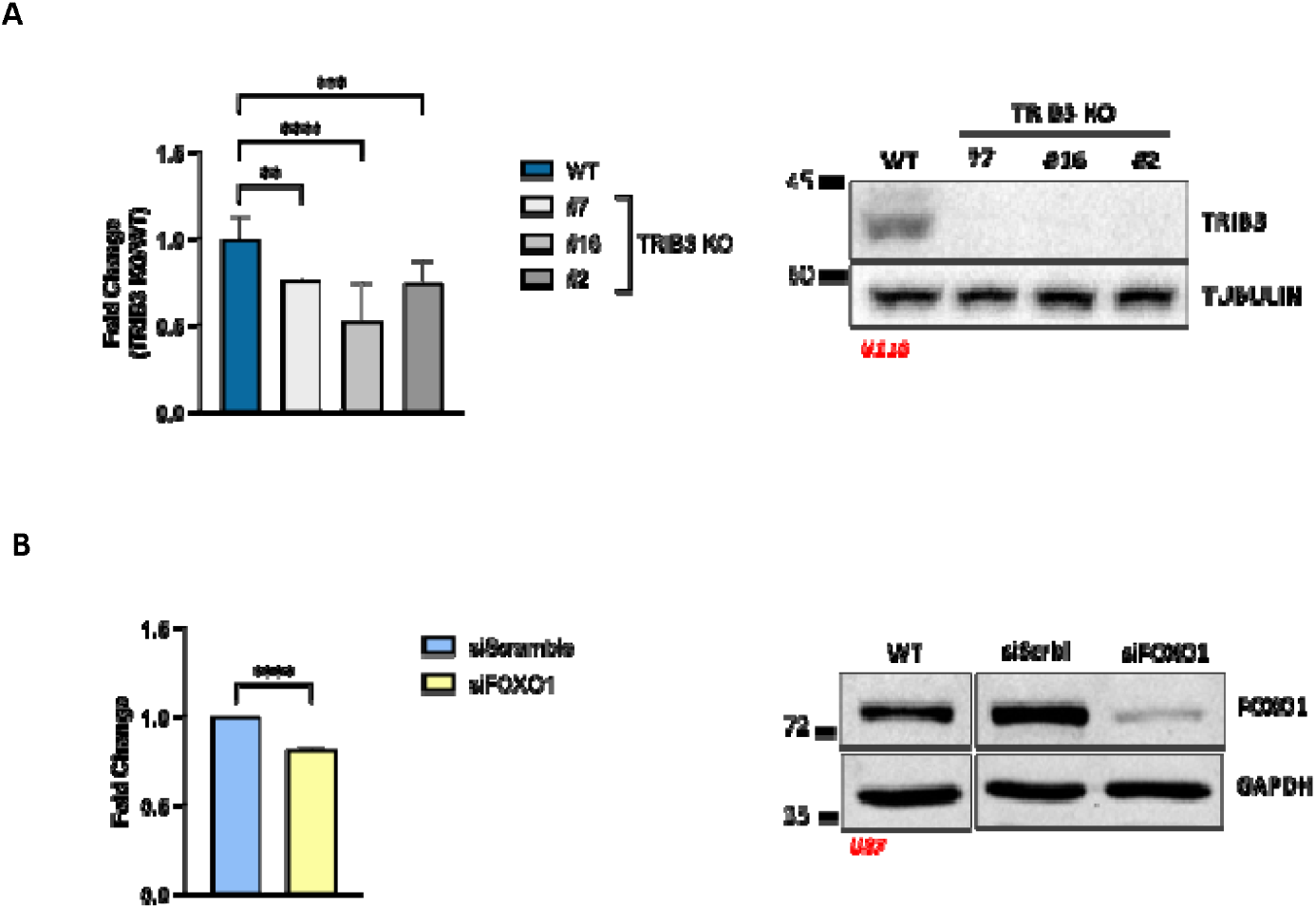
FOXO1 and TRIB3 depletion reduces glioblastoma cell viability. **A:** Cell viability in U118 wild-type (WT) cells compared with three independent TRIB3 knockout (KO) clones. Immunoblot analysis confirms efficient TRIB3 depletion. Data were analyzed using one-way ANOVA. Significance: \*\**p*= 0.0012, \*\*\**p*= 0.0006, \*\*\*\**p*< 0.0001. **B:** Cell viability in U87 glioblastoma cells transfected with FOXO1-targeting siRNA or control scramble siRNA. Immunoblot verifies downregulation of FOXO1 expression. Data was analyzed using an unpaired t-test. Significance. \*\*\*\**p*< 0.0001. Data represent mean ± SEM from three independent experiments (n = 3), with each condition assayed in triplicate wells. Together, these data indicate that both TRIB3 and FOXO1 support glioblastoma cell survival.

Taken together, these results underscore the clinical relevance of the FOXO/Tribbles signature in selecting patients for potential targeted therapies. Notably, 70% of grade 4 gliomas exhibit either a high FOXO1 and low FOXO3/4 signature, elevated TRIB1/2/3 levels, or a combination of both. This highlights the potential of FOXO/Tribbles signatures as an effective tool for guiding treatment strategies.

## Discussion

GB are the most lethal primary brain tumors and are associated with poor response to treatments.^3^ A better understanding of the mechanisms responsible for the progression of the disease revealing therapeutic targets and biomarkers capable of stratifying patients are of utmost importance to improve the clinical outcome. In this study, we identified a prognostic gene expression signature of members of the FOXO and Tribbles protein family members capable of predicting the survival of glioma patients. Dysregulation of both FOXO^43^ and Tribbles^44–46^ family members has been implicated in various cancers, including GB, although their precise roles are highly context-dependent.^47,48^ Thus, dissecting the expression and function of individual isoforms in a tumor-specific context is critical for understanding their role in tumor biology and for uncovering novel avenues for therapeutic intervention. To our knowledge, this study is the first to comprehensively evaluate the simultaneous contribution of all FOXO isoforms (FOXO1, FOXO3, FOXO4, and FOXO6) and all Tribbles isoforms (TRIB1, TRIB2, and TRIB3) in gliomas.

We analyzed four independent datasets: two RNA-seq datasets, one from TCGA^49^ containing 698 samples and the other from CGGA^50^ with 1018 samples of glioma tumors, and two microarray datasets, the REMBRANDT^51^ with 580 samples and the Gravendeel^52^ with 284. We used the TCGA datasets to discover a transcriptional signature that links poor survival of glioma patients to high expression of the transcription factor FOXO1 alongside low expression of FOXO3 and FOXO4, combined with high expression of the pseudokinases TRIB1, TRIB2 and TRIB3. This data is in line with the characterization of FOXO proteins as context dependent tumor suppressor proteins^11^ and studies reporting Tribbles proteins as oncogenic proteins^26^. Validation efforts using CGGA, REMBRANDT and Gravendeel data revealed the compelling association of Tribbles expression with poor clinical outcome and a dramatic effect of FOXO1 and FOXO4 but did not replicate the effect of FOXO3, FOXO6 and TRIB1, in every dataset. These inconsistencies may be attributed to ethnic and genetic differences among the patient cohorts: TCGA and REMBRANDT consist primarily of U.S.-based populations, CGGA represents a Chinese cohort, and the Gravendeel dataset includes European patients.^53,54^ Variability in the range of expression levels for each isoform across datasets can influence patient stratification, leading to significant associations in some cohorts but not in others. Moreover, methodological differences in sample processing and transcriptomic analysis are likely to contribute to the discrepancies. Variations in alignment algorithms, expression quantification techniques, alternative transcript detection, and normalization strategies can all impact the resulting expression profiles.^55–57^ Despite dataset-specific variability, a consistent global pattern emerges across all cohorts: whenever statistical significance is observed, low mRNA levels of FOXO3, FOXO4, and FOXO6, and high expression of FOXO1, TRIB1, TRIB2, and TRIB3, are associated with poorer survival outcomes. Importantly, the data suggests that the combined transcriptional profile of FOXO and TRIB family members holds promise as a biomarker to predict the clinical outcome in gliomas patients. However, further studies are required for population-tailored biomarker validation.

It is important to note that the tumor suppressor function of the members of the FOXO family of proteins is context dependent with significant oncogenic potential under certain circumstances.^14^ Our results support a tumor suppressor role for FOXO3, FOXO4 and FOXO6, while suggesting that FOXO1 may act as an oncogene in gliomas. Notably, consensus from a vast body of research is that FOXO1 functions primarily as a tumor suppressor. However, highlighting the context-dependent nature of FOXO1, emerging evidence indicates it can also exhibit oncogenic functions. Specifically, FOXO1 was found to promote stemness gene expression in GB models, thereby driving tumor progression. Inhibition of FOXO1 in GB cellular models induced apoptosis, further suggesting its oncogenic role in GB.^58^ Additionally, elevated FOXO1 levels have been associated with upregulation of steroid enzymes facilitating androgen synthesis^59^ and increased expression of canonical WNT pathway genes^60^, both of which are linked to enhanced tumor proliferation. However, contrasting studies have reported overexpression of FOXO1 has also been associated with increased sensitivity to TMZ, reduced cell adhesion, and impaired migration and invasion abilities of glioma cells^61^ and FOXO1 inactivation correlated with significantly shorter overall and progression-free survival.^62^ FOXO3 expression has been reported to be markedly downregulated in glioma tissues compared to adjacent normal tissues and has been associated with transcriptional signatures of neuronal differentiation in certain glioma stem cell lines. It also appears to influence glioma stem cell fate by promoting a shift away from quiescence. Restoration of FOXO3 expression induces cell cycle arrest and inhibits glioma cell proliferation and metastasis.^63,64^ Additionally, FOXO3 activation is a key role in enhancing GB sensitivity to EGFR inhibitors.^65^ However, paradoxically, FOXO3 has also been shown to induce resistance to TMZ in glioma cells through distinct molecular pathways.^66,67^ FOXO4 is consistently reported to be downregulated in GB tissues and cell lines relative to normal brain tissue. Its overexpression significantly inhibits glioma cell proliferation, migration, and invasion, while promoting apoptosis *in vitro. In vivo* studies further demonstrate that FOXO4 overexpression suppresses tumor growth.^68^ In our analysis, FOXO4 exhibited the strongest association with patient survival, suggesting a predominant tumor suppressor role in glioma. This effect may be attributed to the relatively higher expression of FOXO4 compared to the other isoforms in supportive cells of the human brain (Fig. 3 C-E). In contrast, FOXO6 displays a more ambiguous functional profile. While its expression is low in adult neural stem cells and not essential for maintaining proliferation, FOXO6 plays a role in mediating cell state transitions, including exit from quiescence and induction of proliferation.^69^ Among all FOXO isoforms evaluated in our study, FOXO6 exhibited the weakest discriminatory power between high and low expression groups in survival analysis and contributed minimally to the overall prognostic signature (Suppl. Fig. 1A). The dual functionality of FOXO proteins as tumor suppressors or oncogenes depending on the cellular environment is poorly understood but might be due to different structural features of these proteins or their levels of expression in different tissues.^18^ Nonetheless, it highlights the critical importance of cellular context and therapeutic timing when considering FOXO-targeted strategies for glioblastoma treatment.

Consistent with previous studies^70^, our results strongly support the role of the Tribbles proteins TRIB1 and TRIB2 as oncogenic drivers in glioma. Interestingly, TRIB1 expression was not only associated with overall survival in GB patients across multiple cohorts, but it’s *in vitro* overexpression reduces RT- and TMZ-induced apoptosis by activating pro-survival signaling pathways, including ERK and AKT.^44,71^ Similarly, TRIB2 has been implicated in resistance to RT and TMZ in glioma, partly through its association with MAP3K1 expression.^45^ The role of TRIB3 in cancer, including gliomas, is complex and points to a highly context-dependent dual activity. Several studies have reported a tumor-suppressor function for TRIB3.^72–74^ Notably, Salazar et al. reported that its upregulation is part of a stress response that promotes autophagy in glioma cells after cannabinoid treatment.^75^ Conversely, other studies have suggested that TRIB3 functions primarily as an oncogene in various malignancies^76–78^, including gliomas^79,80^. We provide evidence that TRIB3 acts as a tumor promoter in human gliomas and drives growth in GB cells. Nevertheless, the context-dependent nature of TRIB3 function is still debated and must be explicitly addressed through future in vivo and clinical research.

Taken together, these observations, along with our data suggest that FOXO1, TRIB1, TRIB2 and TRIB3 may act are drivers of glioma progression, while the loss of FOXO3, FOXO4 and FOXO6 could be contributing to glioma development through tumor-suppressive mechanisms. Therefore, FOXO and Tribbles families should be considered as potential therapeutic targets for this disease. Although non-liganded transcription factors such as FOXO proteins and the pseudokinases of the Tribbles family have been considered undruggable, recent progress provides a variety of strategies to modulate the activity of these proteins.^81^ In particular, several compounds have been identified capable of increasing FOXO activity, including Selinexor ^82^, Dactolisib ^83^ and FOXO4-DRI ^83^. Carbenoxolone ^84^ and AS1842856 ^57^have been reported to interfere with pan-FOXO and specific FOXO1 activity, respectively. Despite these advances, isoform-specific FOXO activators have not yet been identified.^84^ Furthermore, Tribbles modulating agents including TRIB2 inhibitors are emerging.^24,38,85–87^ Hypothetically, a therapeutic strategy combining global FOXO activation and global Tribbles inhibition combined with FOXO1-specific inhibition could help counteract the transcriptional profile associated with high-grade glioma. However, further studies are required to explore the feasibility, efficacy, and safety of such an approach.

In summary, we provide evidence for a potent tumor suppressor role of FOXO4 and a significant oncogenic action of FOXO1 and Tribbles protein family in gliomas. To a lesser degree, the tumor suppressor role of FOXO3 and FOXO6, and oncogenic of TRIB1 might be important for predicting gliomas survival. The FOXO/Tribbles signature can be helpful to stratify patients with glioma predicting the clinical outcomes. An improvement of 4.5 months in medium survival not only is significant but can be meaningful for these patients. We propose to explore the therapeutic potential of FOXO1, FOXO4, TRIB2 and TRIB3 as drug targets to improve the treatment of patients with glioma. Drugs targeting these novel targets, combined with the FOXO/Tribbles signature as companion biomarkers, hold the potential to enhance GB treatment outcomes while sparing patients from ineffective therapies. TRIB2 has been shown to be druggable by approved kinase inhibitors and in the last years several companies have emerged to develop FOXOs therapeutics. This suggests that our results can have an immediate impact by establishing new potential therapeutic targets for several cancer types, in particular gliomas. However, the clinical relevance of these observations and the mechanistic of these proteins in gliomas has yet to be fully established.

## Authorship statement

Conceptualization and methodology: A.T.M., B.I.F. and W.L.; Formal analysis: B.S. and A.B.D; Data acquisition: B.S., A.B.D and A.T.M.; Experimental validation and data analysis: I.G and B.I.F. Visualization: B.S., A.T.M. and B.I.F.; Writing—original draft: B.S., B.I.F. and W.L.; Writing—review & editing: B.S., B.I.F., JMS and W.L.; Supervision: A.T.M., B.I.F. and W.L.; Funding acquisition: B.I.F., A.T.M. and W.L.

## Data availability

The datasets analyzed in this study are publicly available. TCGA RNA-seq data were obtained from The Cancer Genome Atlas (https://portal.gdc.cancer.gov/) and reclassified according to WHO CNS5 criteria. CGGA RNA-seq datasets were downloaded from the Chinese Glioma Genome Atlas (http://www.cgga.org.cn/). Microarray datasets were retrieved from the Gene Expression Omnibus (GEO) under the accession numbers GSE68848 (REMBRANDT) and GSE16011 (Gravendeel). The code and processed data used for the survival analyses are available at: https://github.com/maialab/enduring.

---

**Suppl. Fig. 1**| **High FOXO1, low FOXO3/FOXO4, and elevated TRIB1/2/3 mRNA levels form a molecular signature predictive of clinical outcomes. A:** Kaplan-Meier plot of all possible combinations of low and high levels of FOXOs for TCGA glioma samples. Median value was set as the cut-off Profiles with worse survival are highlighted in red. **B-E:** Kaplan-Meier plots of FOXO for CGGA glioma samples. The lower quartile, in blue, and the upper quartile, in red, were established as the cut-off for low and high expression levels, respectively. **B:** FOXO1; **C:** FOXO3; **D:** FOXO4; **E:** FOXO6. **F:** Kaplan-Meier plot of all possible combinations of low and high levels of Tribbles levels for TCGA glioma samples. Median value was set as the cut-off. Profile with worse survival is highlighted in red. **G-I:** Kaplan-Meier plots of Tribbles for CGGA glioma samples. The lower quartile, in blue, and the upper quartile, in red, were established as the cut-off for low and high expression levels, respectively. **G:** TRIB1; **H:** TRIB2; **I:** TRIB3. **J:** Kaplan-Meier plot of a combination of high levels of FOXO1 and low levels of FOXO3/4 plus high levels of Tribbles for CGGA glioma samples. The median value was set as the cut-off. “Else” means every other possible combination of expressions than the stated in comparison. High FOXO1 and low FOXO3/4 mRNA levels combined with high levels of TRIB1/2/3 (↑FOXO1/↓FOXO3/4/↑TRIBs) are represented in pink. All p-values refer to log-rank test.

**Suppl. Fig. 2**| Kaplan-Meier plot of the different types of tumors for TCGA glioma samples. p-value refers to log-rank test.

**Suppl. Fig. 3**| **A:** Kaplan-Meier plot of a combination of FOXOs and Tribbles levels in TCGA GB samples. **B:** Kaplan-Meier plot of a combination of FOXOs and Tribbles levels in TCGA grade 4 astrocytoma samples. Subgroups with high FOXO1 and low FOXO3/4 mRNA levels (↑FOXO1/↓FOXO3/4/else), high levels of TRIB1/2/3 (else/↑TRIBs), or both (↑FOXO1/↓FOXO3/4/↑TRIBs) represented together in pink. “Else” means every other possible combination of expressions than the stated in comparison. All p-values refer to log-rank test.

