## Supplementary material for "Combined expression of FOXO and Tribbles proteins predicts survival of glioma patients": Suppl. Material

**Suppl. Table 1:** Summary of datasets

| Factor | TCGA | CGGA | REMBRANDT | Gravendeel |
|---|---|---|---|---|
| **Total Number** | 705 | 1018 | 552 | 268 |
| | *Median (LQ, UQ)* | *Median (LQ, UQ)* | Median (LQ, UQ) | Median (LQ, UQ) |
| **Age at diagnosis** (years) | 47.0 (35.0, 59.0) | 42.0 (35.0, 51.0) | - | 51.53 (38.1, 61.4) |
| **Follow-up all cases** (months) | 12.5 (5.3, 23.9) | 27.5 (10.8, 66.7) | 21.9 (10.5, 44.5) | 14.6 (6.7, 39.8) |
| **Follow-up still living** (months) | 12,0 (5.2, 22.6) | 70,9 (42,7, 99,6) | 16.0 (6.0, 31.5) | 69.9 (39.8, 85.8) |
| **Vital status** | | | | |
| **Alive** | 431 (61.1%) | 362 (35.6%) | 87 (15%) | 36 (12.7%) |
| **Dead** | 183 (26.0%) | (617 (60.6%) | 363 (62.6%) | 240 (84.5%) |
| **NA** | 91 (12.9%) | 39 (3.83%) | 130 (22.4%) | 8 (2.8%) |
| **WHO classification** | | | | |
| **Astrocytoma, Grade 2** | 113 (16.0%) | 175 (17.2%) | 61 (10.5%) | 13 (4.6%) |
| **Astrocytoma, Grade 3** | 98 (13.9%) | 214 (21.0%) | 53 (9.14%) | 16 (5.6%) |
| **Astrocytoma, Grade 4** | 24 (3.40%) | - | - | - |
| **Oligodendroglioma, Grade 2** | 81 (11.5%) | 112 (11,0%) | 27 (4.7%) | 8 (2.8%) |
| **Oligodendroglioma, Grade 3** | 70 (9.93%) | 9 (0.9%) | 22 (3.8%) | 44 (15.5%) |
| **Mixed, Grade 2** | - | 94 (9.2%) | 4 (0,7%) | 3 (1.1%) |
| **Mixed, Grade 3** | - | 21 (2.1%) | 3 (0.5%) | 25 (8.8%) |
| **Glioblastoma, Grade 4** | 221 (31.3%) | 388 (38.2%) | 190 (32.7%) | 159 (56.0%) |
| **NA** | 98 (13.9%) | 5 (0.5%) | 192 (28,8%) | - |
| **IDH status** | | | | |
| **wt** | 237 (33.6%) | 435 (42.7%) | - | - |
| **mutant** | 430 (61.0%) | 531 (52.2%) | - | - |
| **NA** | 38 (5.4%) | 52 (5.1%) | - | - |
| **1p19q codeletion status** | | | | |
| **non-codel** | 498 (70.6%) | 728 (71.5%) | - | - |
| **codel** | 169 (24.0%) | 212 (20.8%) | - | - |
| **NA** | 38 (5.4%) | 78 (7.66%) | - | - |
| **MGMT status** | | | | |
| **Methylated** | (67.7%) | 472 (46.4%) | - | - |
| **Unmethylated** | (23.3%) | 376 (36.9%) | - | - |
| **NA** | (9.08%) | 170 (16.7%) | - | - |

**Suppl. Table 2:** Hazard Ratio

| Variables | Hazard Ratio | *p* | 95% Confidence Interval | |
|---|---|---|---|---|
| | | | Lower Bound | Upper Bound |
| **Age** (v. young, <48) | | | | |
| middle age (48-64) | 1.690 | 0.049035 | 1.002 | 2.900 |
| elderly (>64) | 5.304 | 3.54E-08 | 2.931 | 9.600 |
| **Gender** (v. male) | | | | |
| female | 1.091 | 0.621516 | 0.772 | 1.500 |
| **IDH status** (v. mutated) | | | | |
| wt | 3.273 | 0.019807 | 1.207 | 8.900 |
| **1p19q status** (v. codel) | | | | |
| non-codel | 14.061 | 0.000067 | 3.834 | 51.600 |
| **MGMT status** (v. methylated) | | | | |
| Unmethylated | 1.186 | 0.415519 | 0.786 | 1.800 |
| **WHO** (v. Oligodendroglioma, Grade 2) | | | | |
| Oligodendroglioma, Grade 3 | 4.072 | 0.018676 | 1.264 | 13.100 |
| Astrocytoma, Grade 2 | 0.186 | 0.000883 | 0.069 | 0.500 |
| Astrocytoma, Grade 3 | 0.347 | 0.029491 | 0.134 | 0.900 |
| Astrocytoma, Grade 4 | NA | NA | | |
| Glioblastoma | NA | NA | | |
| **FOXO/Tribbles status** (v. else/else) | | | | |
| ↑FOXO1/↓FOXO3/4/↑TRIBs | 1.348 | 0.32957 | 0.740 | 2.500 |
| ↑FOXO1/↓FOXO3/4/else | 0.951 | 0.876803 | 0.506 | 1.800 |
| else/↑TRIBs | 0.985 | 0.948892 | 0.624 | 1.600 |
