## Supplementary material for "Combined expression of FOXO and Tribbles proteins predicts survival of glioma patients": Suppl Figures

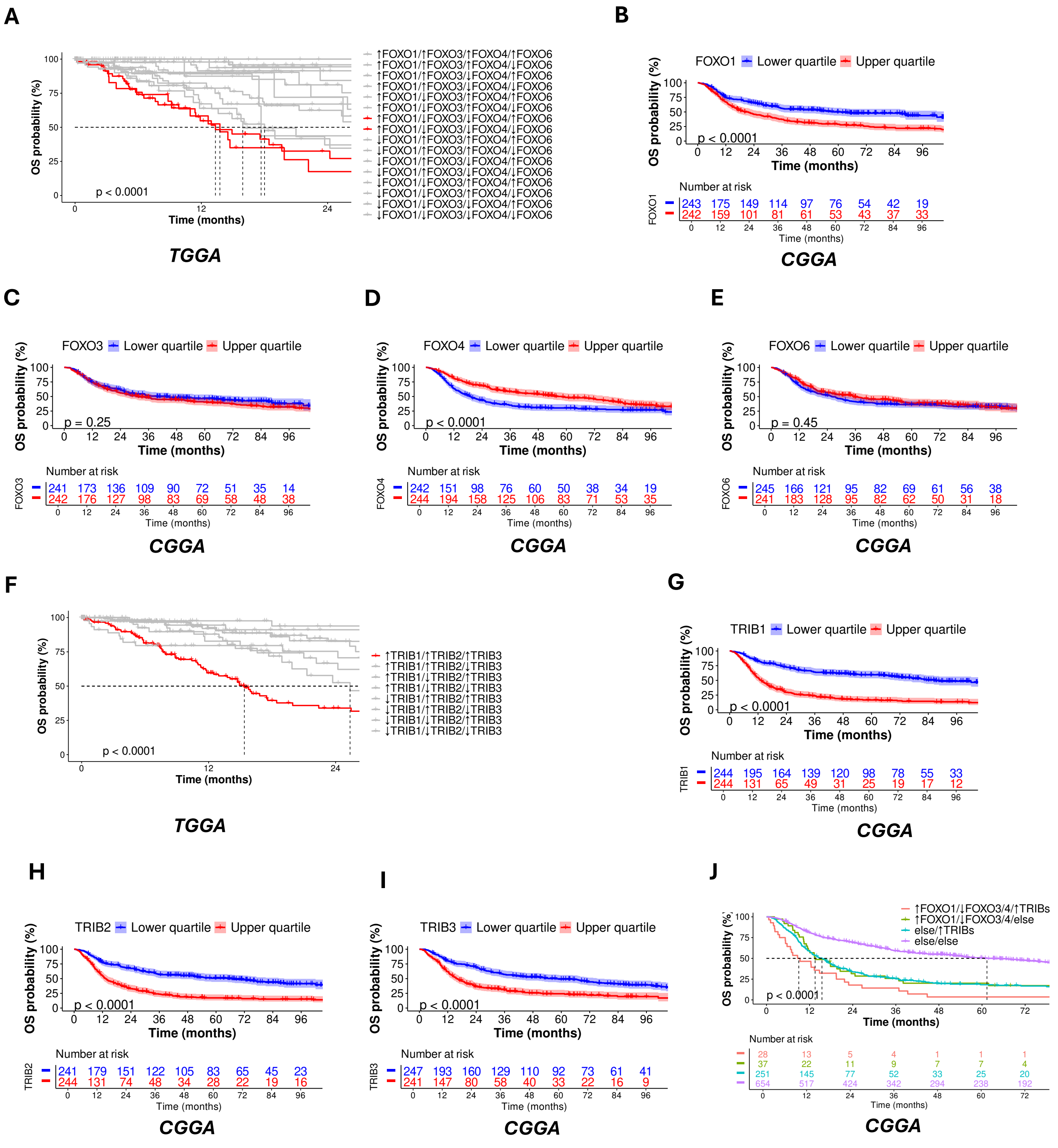


**Fig. Supl. 1| High FOXO1, low FOXO3/FOXO4, and elevated TRIB1/2/3 mRNA levels form a molecular signature predictive of clinical outcomes. A:** Kaplan-Meier plot of all possible combinations of low and high levels of FOXOs for TCGA glioma samples. Median value was set as the cut-off Profiles with worse survival are highlighted in red. **B-E:** Kaplan-Meier plots of FOXO for CGGA glioma samples. The lower quartile, in blue, and the upper quartile, in red, were established as the cut-off for low and high expression levels, respectively. **B:** FOXO1; **C:** FOXO3; **D:** FOXO4; **E:** FOXO6. **F:** Kaplan-Meier plot of all possible combinations of low and high levels of Tribbles levels for TCGA glioma samples. Median value was set as the cut-off. Profile with worse survival is highlighted in red. **G-I:** Kaplan-Meier plots of Tribbles for CGGA glioma samples. The lower quartile, in blue, and the upper quartile, in red, were established as the cut-off for low and high expression levels, respectively. **G:** TRIB1; **H:** TRIB2; **I:** TRIB3. **J:** Kaplan-Meier plot of a combination of high levels of FOXO1 and low levels of FOXO3/4 plus high levels of Tribbles for CGGA glioma samples. The median value was set as the cut-off. "Else" means every other possible combination of expressions than the stated in comparison. High FOXO1 and low FOXO3/4 mRNA levels combined with high levels of TRIB1/2/3 (↑FOXO1/↓FOXO3/4/↑TRIBs) are represented in pink. All p-values refer to log-rank test.

A

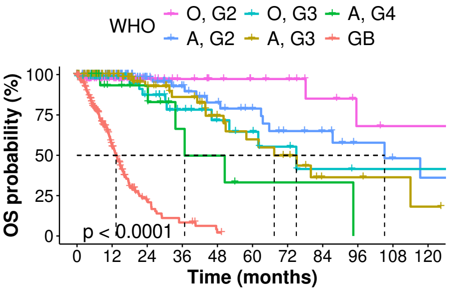


**Fig. Supl. 2|** Kaplan-Meier plot of the different types of tumors for TCGA glioma samples. p-value refers to log-rank test.

A B

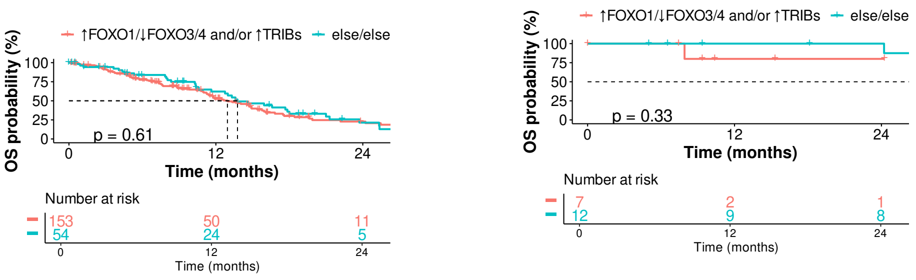


***GB*** ***Astrocytoma, grade 4***

**Suppl. Fig. 3| A:** Kaplan-Meier plot of a combination of FOXOs and Tribbles levels in TCGA GB samples. **B:** Kaplan-Meier plot of a combination of FOXOs and Tribbles levels in TCGA grade 4 astrocytoma samples. Subgroups with high FOXO1 and low FOXO3/4 mRNA levels (↑FOXO1/↓FOXO3/4/else), high levels of TRIB1/2/3 (else/↑TRIBs), or both (↑FOXO1/↓FOXO3/4/↑TRIBs) represented together in pink. "Else" means every other possible combination of expressions than the stated in comparison. All p-values refer to log-rank test.
